# Efficient Functions for Energetic Least-Cost Analysis over Land and Water in the lbmech package for R

**DOI:** 10.1101/2023.07.09.548254

**Authors:** Andrés G. Mejía Ramón

## Abstract

Current least-cost tools for behavioral scientists are computationally insufficient and lack methods to independently derive animal-specific energetic cost functions based on locational data. Moreover, topologically-valid approaches only measure the biomechanical cost of movement, when sometimes net metababolical expediture may be a more-important consideration. In this paper, I first derive generalizable cost functions for energetic expenditure for any arbitrary biped or quadruped, versus the currently-employed regressed functions with limited cross-taxa or behavioral mode applicability. I then describe the cost-distance functions of the R package lbmech, designed to (1) regress the parameters describing behavioral sensitivity to slope from GPS tracking data; and (2) calculate the costs of movement in terms of time, kinetic work, and net metabolic expenditure. Finally, I demonstrate the utility of this package using publicly-available GPS, ocean current velocity, and elevation data from the Balearic Islands of Spain. This approach greatly facilitates the collection of field-based data given the reliance on GPS data. Moreover, it allows us to identify and consider individual- and group-level effects and variances that may lead to differential movement-related fitness with evolutionary consequences.

## 1 Introduction

Wood and Wood (2006) and Berti et al. (2022) have proposed applying least-cost path analysis to an animal’s energetic locomotion expenditure. These studies utilize regressed functions (Pandolf et al. 1977; Pimental and Pandolf 1979; Pontzer 2016) to quantify animals’ net metabolic expenditure. The former conflate mechanical work and energy loss due to basal metabolism and employ a topologically-invalid function (Pandolf et al. 1977). The latter, meanwhile, neglects the energetic cost of being alive over an arbitrary period of time. These distinct processes (cf. Pontzer et al. 2016) — though related— may uniquely influence movement decision-making across landscapes. Mechanical work signifies real-time mechanical effort, while energy integrates both mechanical effort and life’s energetic cost. Additionally, current least-cost tools for behavioral scientists are computationally insufficient and lack methods to independently derive animal-specific energetic cost functions based on locational data.

In least-cost path analyses, typically three cost types are considered for landscape traversal: (1) dimensionless, expressing relative cost (e.g., location A has tenfold travel cost compared to location B); (2) temporal, primarily for human movement based on Tobler’s Equation for average human walking speed per incline; and (3) energetic, using either Pandolf’s Equation for human net metabolic expenditure per incline, or Pontzer’s function for estimating an animal’s metabolic expenditure per incline based on its mass. Each approach has various shortcomings.

Dimensionless costs —-often based on subjective landcover transport considerations — are simple to generate, but generally lack empirical grounding and are often pseudo-quantitative. Temporal costs — easily conceptualized and linked to quantifiable behavior — lack a simple non-human animal slope-velocity model estimation method, while Tobler’s hiking function’s parameter variability remains poorly understood. Net metabolic expenditure combining basal metabolism and kinematic work, can misrepresent energy variables. Notably, Pandolf’s human transport equation cannot predict downhill movement, yielding implausibly negative energy estimates (Herzog 2014), suggesting the possibility of energy gain from downhill travel. Pontzer’s function (2016)—which underlies Berti et al. (2022)’s energetic model and the associated `enerscape` toolkit — derives the cost-of-travel from the biomechanics of muscle contractions. While it has high explanatory power, the large variance — particularly on a logarithmic scale — suggests limited predictive ability in cross-species applications. Thus, a topologically-correct approach that can derive species-specific functions from empirical data is needed.

In this paper, I first derive generalizable cost functions for any arbitrary movement, based on energetics and mechanical physics, not regressed functions with limited cross-taxa or behavioral mode applicability. This allows for three independent locomotive energy expenditure types: (1) kinematic/mechanical work; (2) gravitational kinematic/mechanical work; and (3) basal metabolic expenditure. The second section introduces the `lbmech` package, initially designed for the analyses described herein and to determine energetic parameters for specific taxa and behavioral mode using locational data. Lastly, I illustrate the framework using GPS, topographic, and ocean current data from the *Illes Balears* (Lera et al. 2017).

## 2 The Energetics of Locomotion

To traverse a terrestrial landscape, kinetic work (*K*) and work against gravity (*U*) are performed. Kinetic work for human movement is 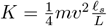 (Kuo 2007), and for quadrupeds *K* = *m*(0.685*v*+0.072) (Heglund et al. 1982). Here, *m* is mass, *v* is speed, *ℓ*_*s*_ is step length, and *L* is leg length. Gravitational work is nonholonomic as it is expended only when moving uphill, resulting in *U* = *mgh*, to lift mass *m* to height *h* with gravity *g*. The total work *W* is:

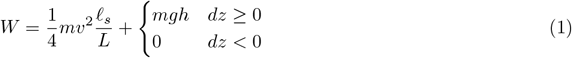

Tobler’s Hiking function predicts velocity *v* based on landscape topography:

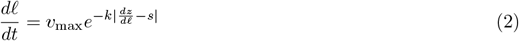

where *ℓ* is horizontal movement, *z* vertical movement, *v*_max_ the maximum speed, *s* the optimal slope, and *k* the speed variation sensitivity. Canonical values for humans are *v*_max_ = 1.5 m/s, *k* = 3.5, and *s* = −0.05. However, `getVelocity()` in the `lbmech` package can extract these variables from GPS data.

Considering a biomechanical inefficiency efficiency factor of *ε*, and adding base metabolic rate 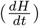, we model the total energy expenditure *E* for bipeds as:

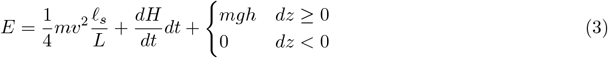

The derived functions are anisotropic due to dependence on 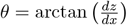, and nonholonomic due to their dependence on gravity. However, conventional least-cost tools such as most GIS toolkits are inadequate given that they operate on cell values themselves. Given that this is a problem of integrating a path integral over a differential surface, alternate methods are required.

A least-cost path algorithm selects paths where segments tend towards the lower end of cost plots in Figure 1. Considering canonical parameters and time minimization (Figure 1a), paths with slopes near the fastest travel slope (*s* = −0.05), are favored. Moving away from this value exponentially increases the cost. For kinetic energy minimization (Figure 1b), extreme slopes are chosen, as movement and kinetic energy are minimal. For work against gravitational potential energy (Figure 1c), any negative slope is equally favored with positive slopes linearly increasing in cost. Downhill movement is primarily kinetic energy-restrained, while uphill movement is mainly limited by gravitational potential energy. For net metabolic energy, penalties apply to slower, mechanically efficient routes due to metabolic costs. Therefore, the most efficient slopes exist at the global minimum of 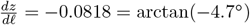 with canonical parameters.

**Figure 1.**
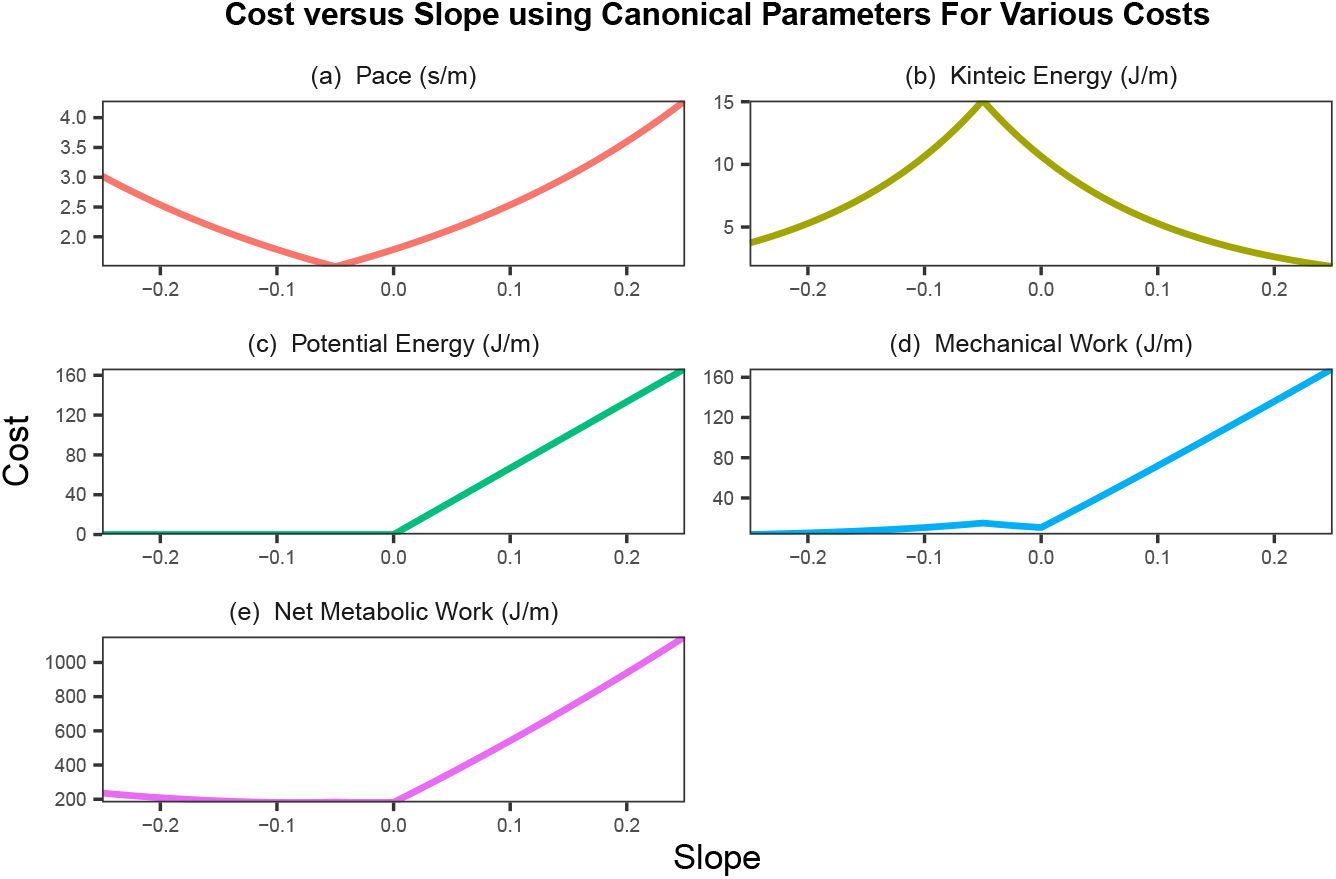
Cost functions using the canonical parameters for Tobler’s hiking function. *k* = 3.5, *s* = −0.05 *m* = 68 kg, *v*_max_ = 1.5 m/s, *L* = 0.75 m, *ℓ*_*s*_ = 1.6 m, *g* = 9.81 m/s^2^, 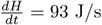, and *ε* = 0.2.

Least-cost toolkits often neglect potential water-based movement—much simpler to calculate than on land. The water surface is represented by a two-dimensional vector field for water surface velocity 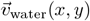. For still water, 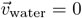. The velocity of someone on water is:

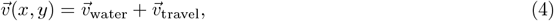

where 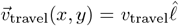 is the travel speed (e.g. swimming or rowing) over still water and 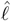 is the unit vector of the intended travel direction. If 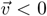, then travel in that direction is impossible. For simplicity, we assume that the caloric cost is proportional to 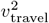.

## 3 The Cost-Distance Functions of lbmech

`lbmech` is a geospatial R package developed for least-cost path analysis, incorporating tools for determining time and energy-based travel costs for animals and humans navigating landscapes. It also includes built-in statistical tools for deriving cost functions from GPS data. Some features, such as estimating potential energetic net primary productivity from tabulated data, are currently under development, and will be discussed in future works.

The general philosophy behind this package is that energetic and least-cost path analyses should always be simple, the most time-taking parts should be done at most once, and ideally that costs should be rooted in empirical reality. It shares features with the `gdistance` library (van Etten 2017), but differs in computation and usability. Both libraries use differential surfaces to represent movement costs between landscape locations and store these costs in ways that enumerate possible transitions. They then perform computations before generating a network `igraph` object for efficient distance calculations (Csardi and Nepusz 2006). However, `lbmech` excels in speed, simplicity, and flexibility due to its data structure and handling methods.

The `gdistance` library uses conductance (i.e., 1/resistance) to store movement costs in a sparse matrix. The use of conductance reduces memory usage but can result in integer overflow errors with large datasets given the need of index masking for when *f*(0) ≠ 0. Meanwhile, `lbmech` stores resistance values directly as an adjacency list and performs linear algebra using the `data.table` package, enabling simpler syntax for many operations. Each possible movement is stored as a unique row, with entries for `‘from’` and `‘to’` nodes, and the difference in or raw final and initial values of the raster. Nodes are named after raster cell coordinates and are stored as `‘x,y’` character strings. This approach facilitates in-place modification of objects, bidirectional raster analysis, and transformations of large rasters without encountering integer overflow limits.

`lbmech` also compartmentalizes the cost-distance workflow, ensuring that intensive steps only need to be performed once. It supports large spatial regions and fine spatial resolutions, critical for land-based transport decision-making at meter and decameter scales. Files are stored in a temporary directory by default, but a workspace directory can be designated for repeated analysis, ensuring efficient file management. Geographic projections and transformations are performed using the `terra` package, while read/write operations involve the `fst` package leveraging speed enabling fast referential and partial access to datasets (particularly on solid state drives). `lbmech` can use multiple elevation rasters, sourced from a polygon grid pointing to a URL or filepath with the region’s elevation data. If no sources are provided, SRTM tiles are fetched using the `elevatr` package.

The package emphasizes empirical reality by focusing on time and energy-based considerations for land-scape movement. Its `energyCosts()` function estimates maximum walking speed at a given slope using a generalized form of Tobler’s hiking function, and one of three biomechanical models (Kuo 2007, Heglund et al. 1982, Pontzer 2016, and an oscillator) for work per unit time or stride. Unlike tools like enerscape for `R`, `lbmech` estimates various types of energetic losses and provides a method to derive cost functions from GPS data through the `getVelocity()` function for use with the Kuo (2007) or Heglund et al. (1982) or oscillator methods. This is less invasive than VO_2_ meters used for net energetic expenditure estimation. Finally, `lbmech` incorporates the `geosphere` package’s distance functions, allowing geodesic corrections at all calculation stages.

## 4 Case Study — Movement around the Balearic Islands

The full code, which includes all functions, figures, and case study, is available on the `lbmech` website. This section outlines the general usage of the package using an example from the Balearic Islands, Spain. The package can be installed within `R`:

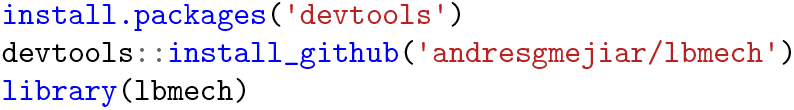

### 4.1 Estimating Velocity

Our GPS dataset of 15,296 usable GPX tracks from https://wikiloc.com, was first detailed by Lera et al. (2017), who analyzed hiking trail activity and network organization seasonality. Given a directory of `.gpx` files, tracks can be imported as a data.table using `importGPX()`:

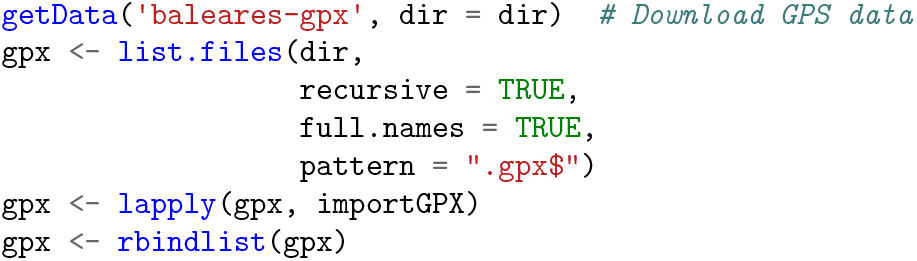

GPS tracks can provide a per-length sample rate finer than even the smallest raster pixels we might use, so they are downsampled to an equivalent rate—approximately a pixel length per GPS sample. For an expected pixel size of 50 m and a maximum speed close to 1.5 m/s, `t_step` = 50/1.5. The `downsampleXYZ()` function is applied for this purpose:

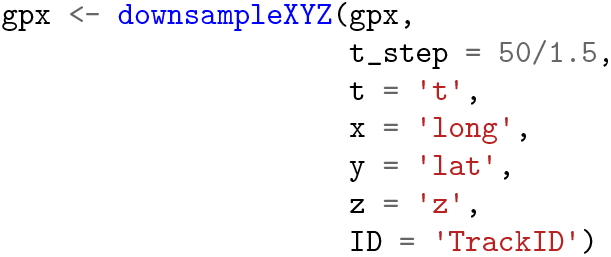

Subsequently, the xyz data is input into `getVelocity()`, which for each GPS point sequence (1) calculates the elevation changes 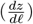 and planimetric speed 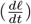, and (2) carries out a nonlinear quantile regression to achieve a Tobler-like function. This results in a list comprised by the nonlinear model, model parameters, and the transformed data.

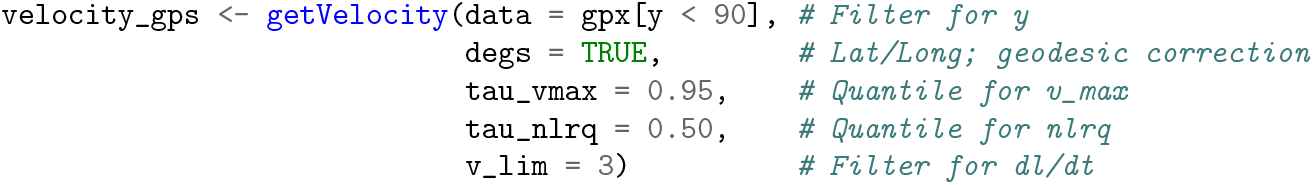

`plotVelocity()` can be used to plot the log-transformed probability of a given speed at a given slope versus the regressed function, as in Figure 2:

**Figure 2.**
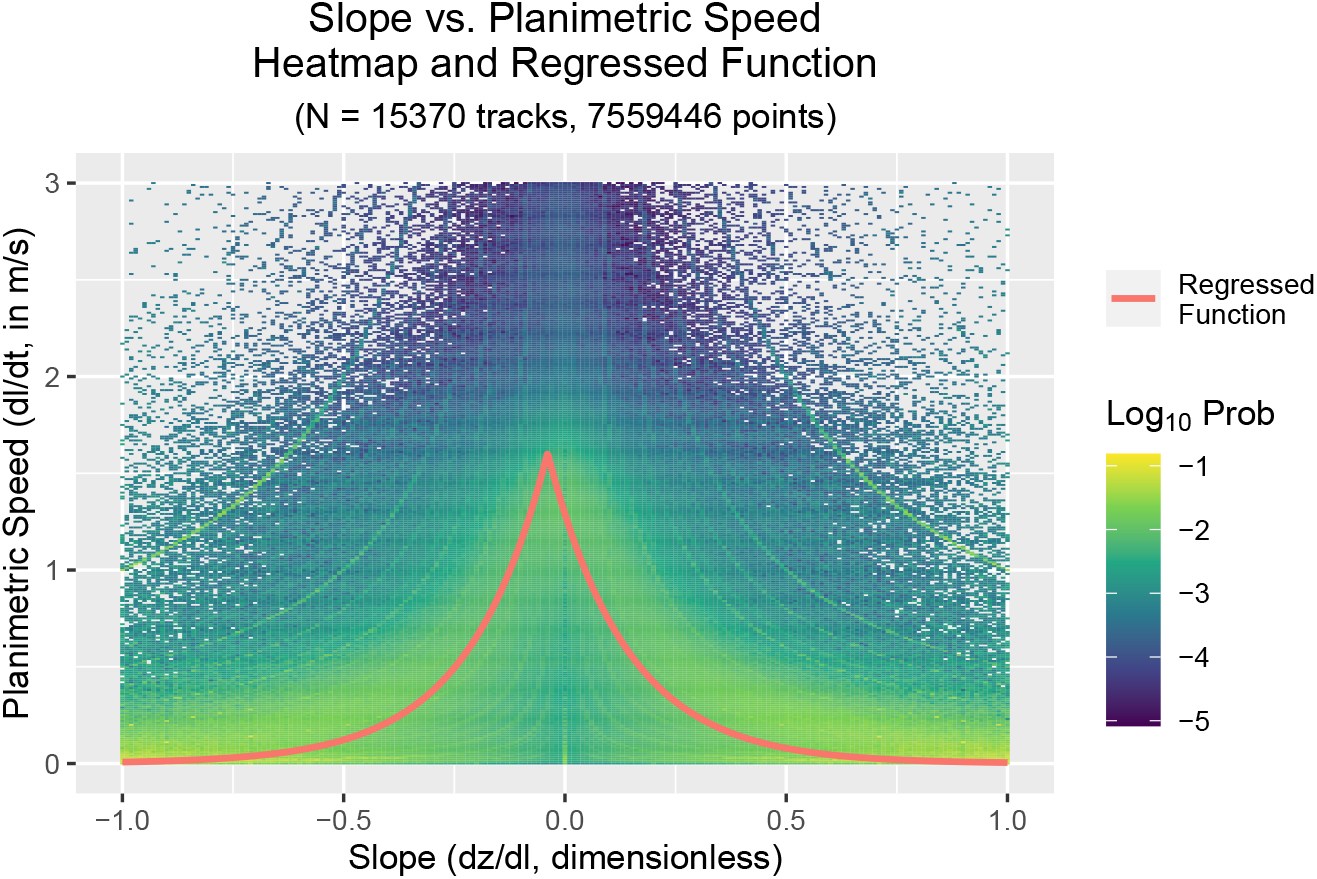
Log-transformed probability of movement at a given speed versus slope for the data reported by Lera et al. (2017).

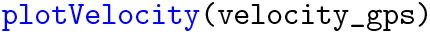

### 4.2 Energetic Cost-Distance Analysis

For efficient data handling in various stages of the workflow — particularly during model-building — a consistent directory should be set.

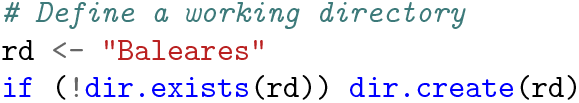

`lbmech` offers automatic terrain data download. Nevertheless, to incorporate ocean movement, we import ocean currents around Mallorca and Menorca — the principal Balearic Islands — on June 13, 2022 previously downloaded from https://data.marine.copernicus.eu/. Subsequently, the maximum movement region (the “world”) will be defined around the raster’s extent:

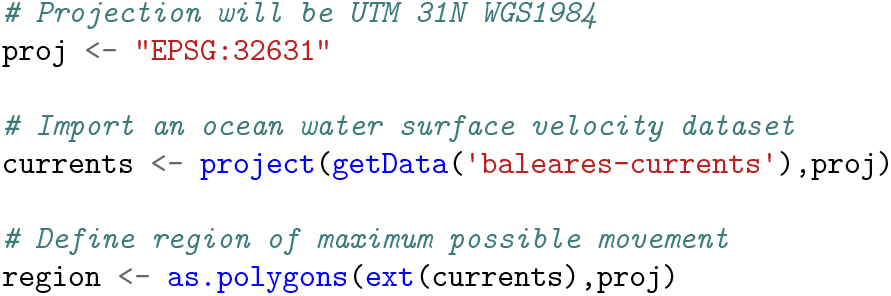

To conduct raster operations, we must define a consistent projection and grid. Unlike most raster packages, `lbmech` requires only the resolution and offsets, not the spatial extents. This can be done with the `fix_z()` function:

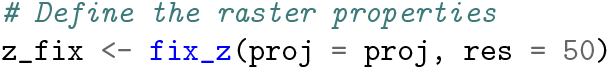

Often, we define a ‘world’ coincident with a digital elevation model. `makeGrid()` can generate a polygon from a raster or filepath. This function helps download SRTM data as needed.

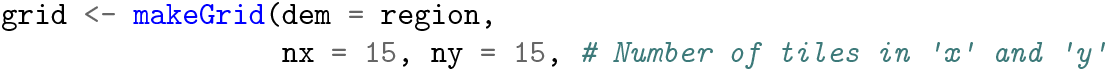

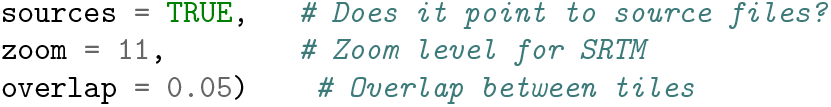

Similar polygons pointing to elevation data sources are often distributed by state GIS agencies. The `defineWorld()` function segments the movement world into manageable, overlapping ‘sectors’, read-in only as required due to memory limitations. This function mainly prepares the data for calculations.

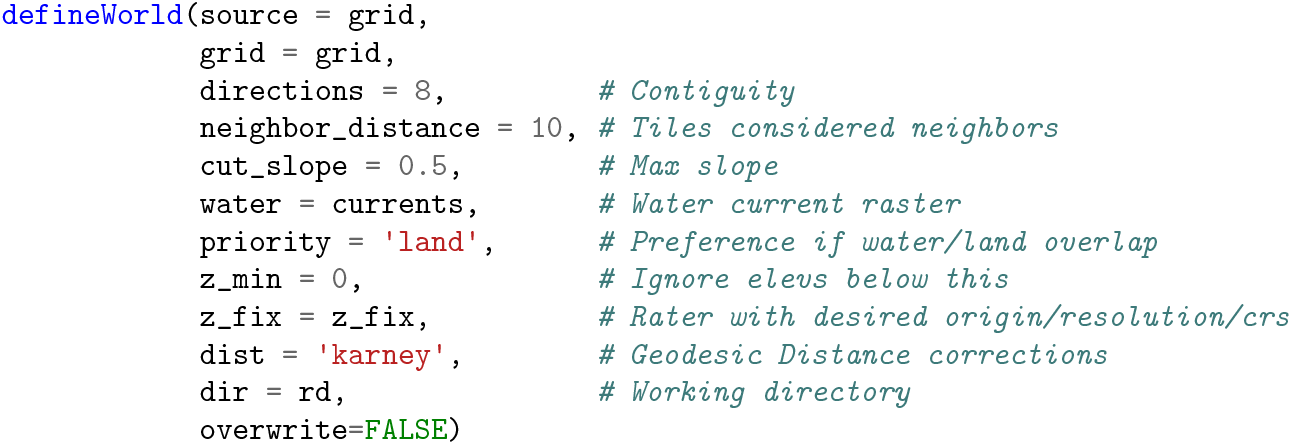

Once a world has been defined (Figure 3), a cost function can be applied using `calculateCosts()`. `energyCosts()` is provided to perform least-time, least-work, and least-energy analyses, as previously described. It prepares data for calculations, but doesn’t typically perform calculations itself.

**Figure 3.**
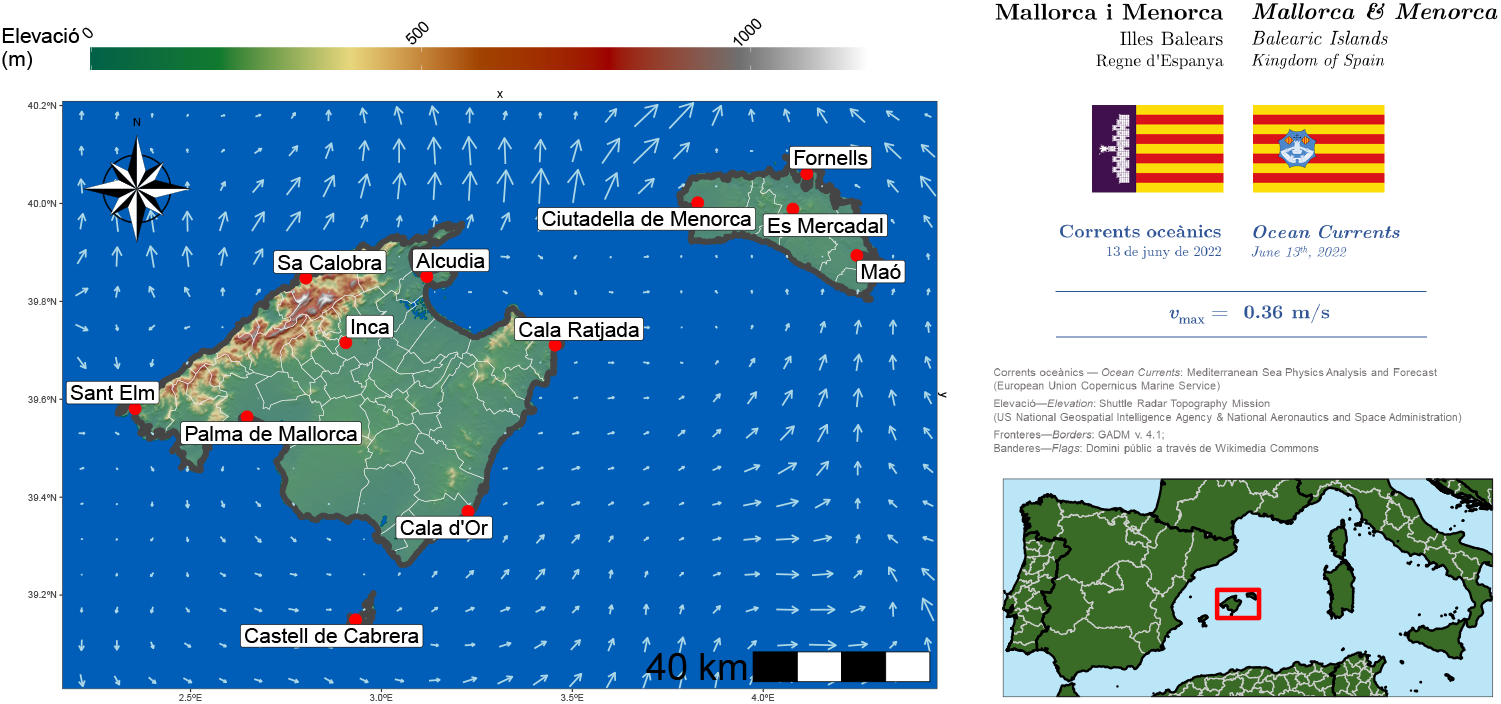
Topography of and currents around Mallorca, Menorca, and Cabrera on June 13^th^, 2022.

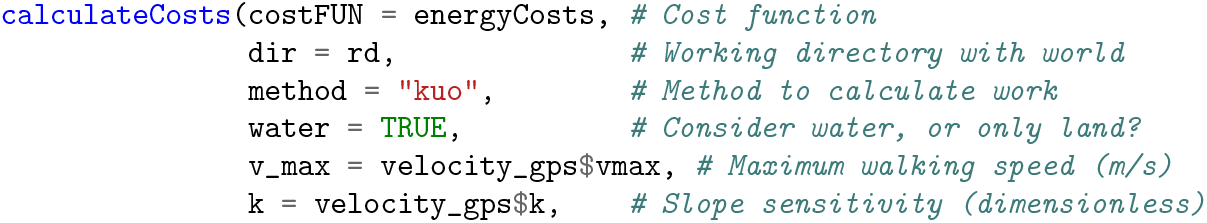
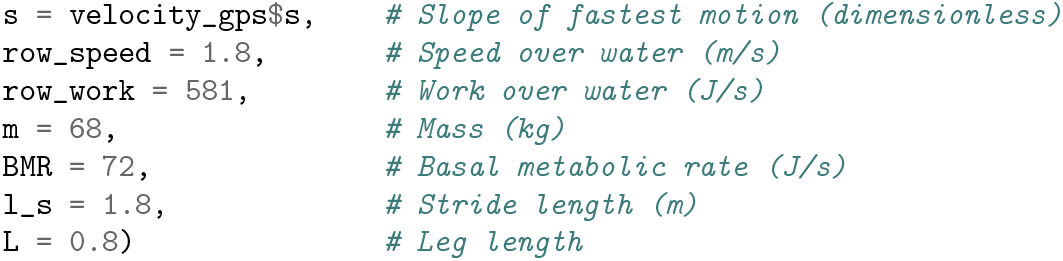

For a given cost function and set of origins/destinations, `getCosts()` identifies necessary sectors, checks and downloads data, performs calculations, and saves cell-wise transition cost tables to the working directory. This way, they won’t need to be fetched or preprocessed again for future calculations.

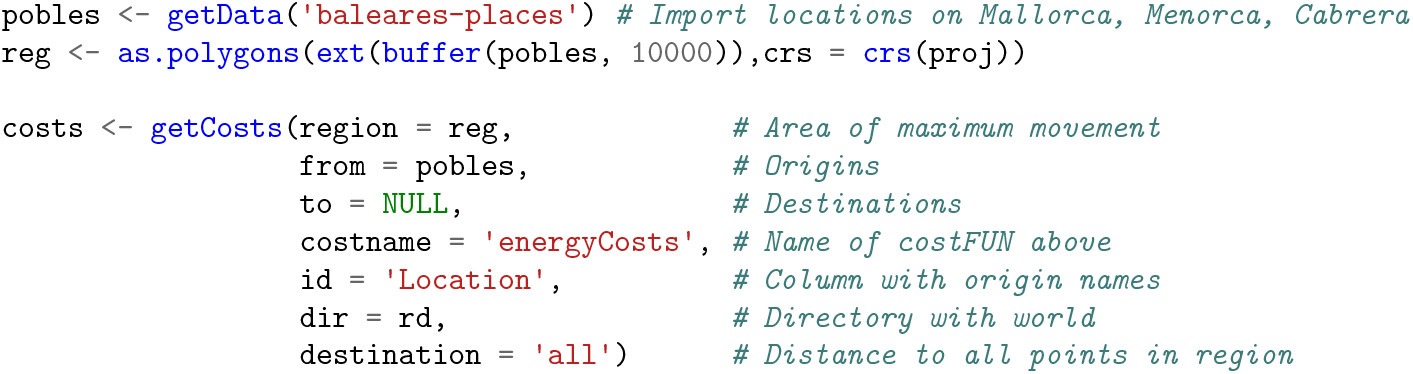

Least-cost paths can be computed for a given set of nodes, here between Palma de Mallorca and Maö, or in Figure 4 :

**Figure 4.**
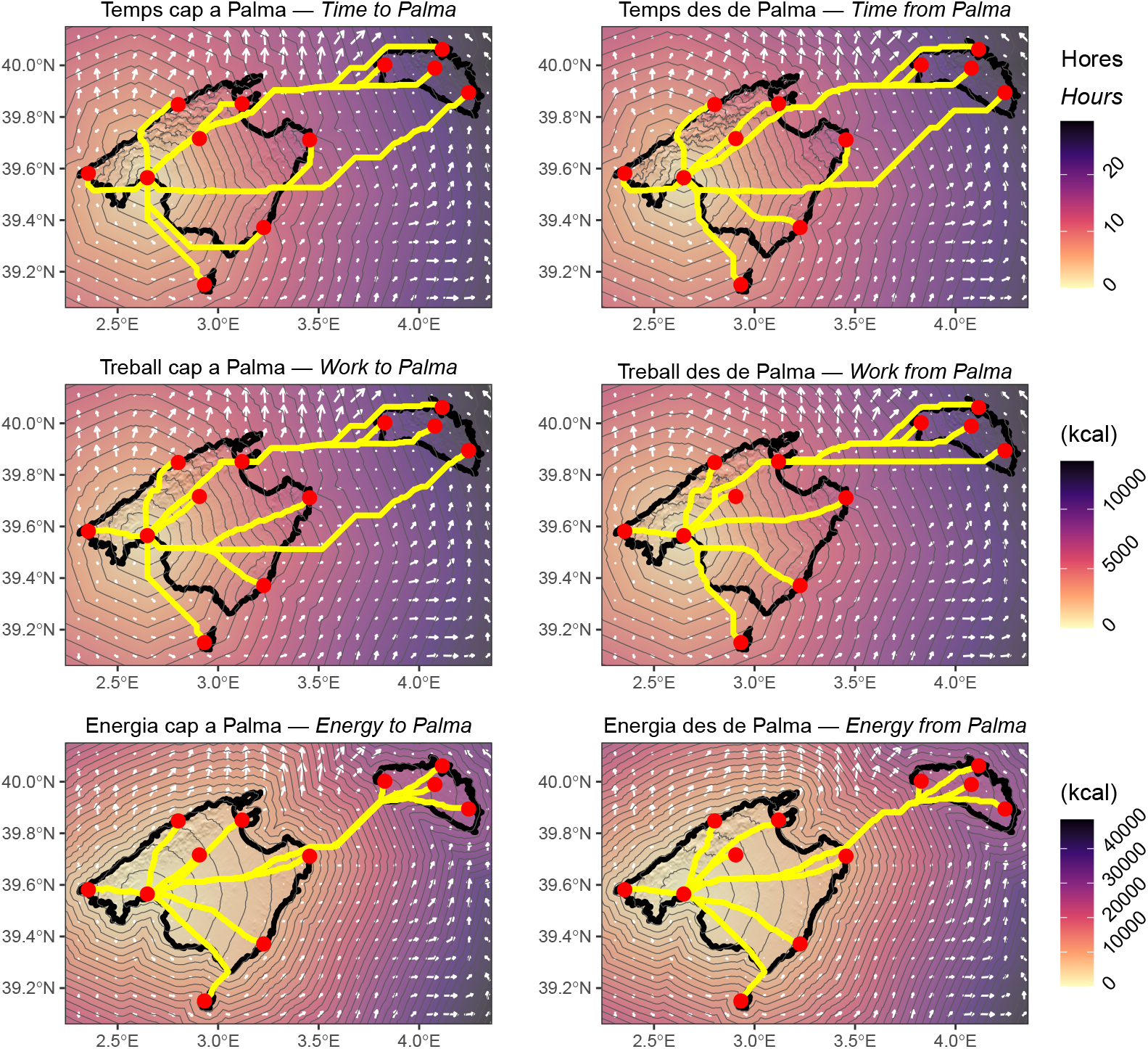
Least-cost paths to and from Palma de Mallorca.

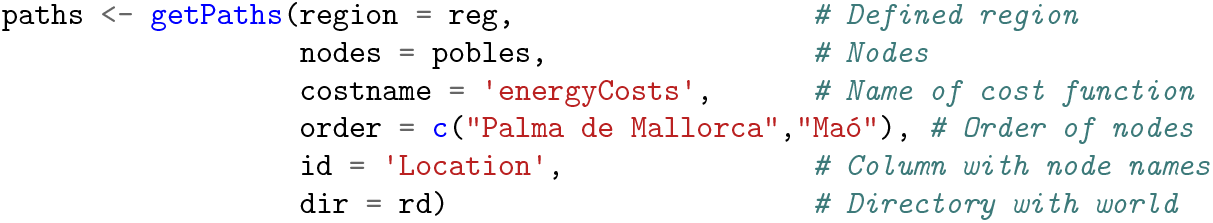

Finally, corridors specify the minimum detour cost to a particular location on a path between two or more points, relative to the least cost path between those points. This can be calculated from the output cost rasters utilizing the `makeCorridor()` function, here from Castell de Cabrera to Maö via Sa Calobra:

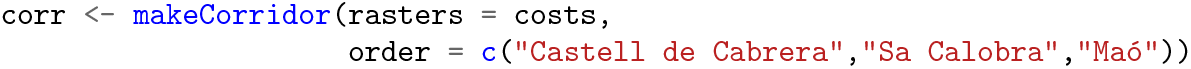

## 5 Conclusion

This method improves on cost-distance analysis in behavioral and spatial sciences by modeling individual- and group-level effects, tailoring empirical costs, and efficiently handling large-scale, high-resolution data. For instance, we regressed unique parameters for human movement in the Balearic Islands using GPS data, facilitating nuanced behavioral response analyses. Moreover, `lbmech` offers significant computational benefits over standard methods, intuitively combining water and land transportation while processing extensive regions at 50m resolution, all without overflow errors on a 32 GB laptop. Segmentation of GPS tracks by individual or population would allow us to potentially identify differences in behavioral responses to landscape slope and their energetic consequences. This has direct implications with regards to differential fitness, opening the possibility for evolutionary studies of locomotive efficiency in specific landscapes and the estimation of the minimum amount of energy required to sustain mobile behavior.

## Supporting information

Inline Code & Figures

## Data Availability Statement

The code for all analyses can be obtained at: https://github.com/andresgmejiar/lbmech

## Acknowledgments

The author would like to thank W. David Walter, Douglas Miller, Douglas Bird, Mary Shenk, Christian John, Kenneth Hirth, Tom Froese and Georgii Karelin for comments and suggestions on previous versions of the model.

The author would like to thank W. David Walter, Douglas Miller, Douglas Bird, Mary Shenk, Christian John, Kenneth Hirth, Tom Froese and Georgii Karelin for comments and suggestions on previous versions of the model. Elio González kindly guided the author through a number of the trails, for which he is hugely grateful.

## Notes

https://github.com/andresgmejiar/lbmech

### Competing Interest Statement

The authors have declared no competing interest.

