## Supplementary material for "Efficient Functions for Energetic Least-Cost Analysis over Land and Water in the lbmech package for R": Inline Code & Figures: movement-raw.RMD

````
---
title: "Raw Cost Analysis Code"
author: "Andrés G. Mejía Ramón"
date: "June 6, 2023"
md_document:
variant: markdown_github
github_document:
toc: true
pandoc_args: --webtex "https://latex.codecogs.com/svg.latex?"
---

# Preparation

The package can be installed within R:

```{r setup}  
#install.packages('remotes')
remotes::install_github('peer-review-53297/lbmech-peer')
library(lbmech-peer)
dir <- getwd()
```

# Velocity Estimation

The GPS dataset of 15,296 usable GPX tracks from [https://wikiloc.com](https://wikiloc.com) was first detailed by Lera et al.
(2017), who analyzed hiking trail activity and network organization seasonality. Given a directory of .gpx
files, tracks can be imported as a data.table using `importGPX()`:

```{r import}
getData('baleares-gps', dir = dir) 
gpx <- list.files(dir ,
recursive = TRUE,
full.names = TRUE,
pattern = ".gpx$")
gpx <- lapply(gpx, importGPX)
gpx <- rbindlist(gpx)
```

Given that GPS tracks can provide a per-length sample rate finer than even the smallest raster pixels we
might use, these tracks are downsampled to an equivalent rate—approximately a pixel length per GPS
sample. For an expected pixel size of 50 m and a maximum speed close to 1.5 m/s, `t_step` = 50/1.5. The
`downsampleXYZ()` function is applied for this purpose:

```{r downsample}
gpx <- downsampleXYZ(gpx, t_step = 50/1.5,
t = 't', x = 'long', y = 'lat', z = 'z', 
ID = 'TrackID')
```

Subsequently, the xyz data is input into getVelocity(), which for each GPS point sequence (1) calculates
the elevation changes (dz/dl) and planimetric speed (dl/dt), and (2) carries out a nonlinear quantile regression to
achieve a Tobler-like function. This results in a list comprising the nonlinear model, model parameters, and
the transformed data.

```{r getVelocity}
velocity_gps <- getVelocity(data = gpx[y < 90], # Filter for y
degs = TRUE,        # Lat/Long; geodesic correction
tau_vmax = 0.95,    # Quantile for v_max
tau_nlrq = 0.50,    # Quantile for nlrq
v_lim = 3)          # Filter for dl/dt

print(velocity_gps[1:(length(velocity_gps)-1)])

plotVelocity(velocity_gps)
```

# Cost-Distance Analysis

For efficient data handling in various stages of the workflow, particularly during model-building, a consistent
directory should be set.

```{r workingDir}
# Define a working directory
rd <- "Baleares"
rd <- paste0(dir,"/",rd)
if (!dir.exists(rd)){
dir.create(rd)
}
```

`lbmech` offers automatic terrain data download. Nevertheless, to incorporate ocean movement, we also need to import ocean currents around Mallorca and Menorca---the principal Balearic Islands on June 13, 2022---previously downloaded from \url{https://data.marine.copernicus.eu/}. Subsequently, the maximum movement region (the 'world') is defined around the raster's extent:

```{r currents}
# Projection will be UTM 31N WGS1984
proj <- "EPSG:32631"

# Import an ocean water surface velocity dataset
currents <- project(getData('baleares-currents'),proj)

# Define region of maximum possible movement
region <- as.polygons(ext(currents),proj)

# Make ID Column
region$ID <- 1
```

To conduct raster operations, we must define a consistent projection and grid. Unlike most raster packages, lbmech requires only the resolution and offsets, not the spatial extents. This can be done with the fix_z()
function:

```{r z_fix}
# Define the raster properties
z_fix <- fix_z(proj = proj, 
             res = 50)
```

Although a 'world' can be defined directly based on a digital elevation model, generally it is easier to define a polygon coincident with the coverage of a digital elevation model and an attribute pointing to a URL/filepath with the source. The `lbmech::makeGrid()` can make such a polygon for a raster or filepath input, while using a polygon will induce future functions to download SRTM data as-needed. Similar polygons are frequently distributed by state GIS agencies:

```{r makeGrid}
# Make a grid
grid <- makeGrid(dem = region,     # Input; here polygon for SRTM
               nx = 15, ny = 15, # Cols/Rows to divide the polygon
               sources = TRUE,   # Just crop/divide, or point to source?
               zoom = 11,        # Zoom level for SRTM data 
               overlap = 0.05)   # Fraction overlap between adjacent tiles
```

The `defineWorld()` function segments the movement world into manageable, overlapping `sectors', read-in only as required due to memory limitations:

```{r defineWorld}
# Define world of motion within a workspace
defineWorld(source = grid,            # Elevation source, like makeGrid() output
grid = grid,              # How to divide up the world
directions = 8,           # How adjacency between cells are defined 
neighbor_distance = 10,   # Overlap between tiles in addition to ^
cut_slope = 0.5,          # Maximum traversible slope
water = currents,         # Water velocity source
priority = 'land',        # If data for land or water, who wins
z_min = 0,                # Minimum elevation, below NA in land
z_fix = z_fix,            # Grid with defined projection, resolution
dist = 'karney',          # Geodesic correction method
dir = rd,                 # Working directory
overwrite=FALSE)
```

Once a world has been defined, a cost function can be applied using `calculateCosts()`. `energyCosts()` is provided to perform least-time, least-work, and least-energy analyses, as previously described. It prepares data for calculations, but doesn't typically perform calculations itself.

```{r calculateCosts}
calculateCosts(costFUN = energyCosts,  
             dir = rd,                    # Working directory with world
             method = "kuo",              # Method to calculate work
             water = TRUE,                # Consider water, or only land?
             v_max = 1.6,   # Max walking speed (m/s)
             k = 5.2,          # Slope sensitivity (dimensionless)
             s = 0.04,          # Slope of fastest motion (dimlss.)
             row_speed = 1.8,             # Speed over water (m/s)
             row_work = 581,              # work over water (J/s)
             m = 68,                      # Mass (kg)
             BMR = 72,                    # Basal metabolic rate (J/s)
             l_s = 1.8,                   # Stride length (m)
             L = 0.8)                     # Leg length
```

For a given cost function and set of origins/destinations, `getCosts()` identifies necessary sectors, checks and downloads data, performs calculations, and saves cell-wise transition cost tables to the working directory. This way, they won't need to be fetched or preprocessed again for future calculations.

```{r getCosts}
# Import locations on Mallorca, Menorca, and Cabrera
pobles <- getData('baleares-places')
pobles <- project(pobles,proj)
reg <- as.polygons(ext(buffer(pobles, 10000)),crs = crs(proj))

costs <- getCosts(region = reg,              # Area of maximum movement
from = pobles,             # Origins
to = NULL,                 # Destinations 
costname = 'energyCosts',  # Name of costFUN above
id = 'Location',           # Column with origin names
dir = rd,                  # Directory with world
destination = 'all')       # Distance to all points in region
```

Least-cost paths can be computed for a given set of nodes, here between Palma de Mallorca and Maó:

```{r getPaths}
# Find all least-cost paths between Palma de Mallorca and Maó

paths <- getPaths(region = reg,
nodes = pobles,
costname = 'energyCosts',
order = c("Palma de Mallorca","Maó"),
id = 'Location',
dir = rd)

```

Corridors specify the minimum detour cost to a particular location on a path between two or more points, relative to the least cost path between those points. This can be calculated from the output cost rasters utilizing the `makeCorridor()` function, here from Castell de Cabrera to Ma\'o via Sa Calobra:

```{r makeCorridor}
corr <- makeCorridor(rasters = costs, 
order = rev(c("Castell de Cabrera","Sa Calobra","Maó")))
```


# Code for the figures

## Cost functions

```{r costFuns}
library(data.table)
library(ggplot2)
funs <- data.table(Slope = seq(-0.25,0.25,length.out = 1000)
)[, `(a)  Pace (s/m)` := 1.5 * exp(3.5 * abs(Slope + 0.05))
][, `(b)  Kinteic Energy (J/m)` := 1 / 2 *  68 * `(a)  Pace (s/m)`^-2 
][, `(c)  Potential Energy (J/m)` := fifelse(Slope > 0, 68 * 9.81 * Slope, 0),
][, `(d)  Mechanical Work (J/m)` := `(b)  Kinteic Energy (J/m)` + `(c)  Potential Energy (J/m)`
][, `(e)  Net Metabolic Work (J/m)` := `(d)  Mechanical Work (J/m)`/0.2 + 72 * `(a)  Pace (s/m)`]

f <- melt(funs,id.vars = 'Slope', variable.name = 'Cost')

gg <- ggplot(f, aes(x = Slope, y = value, color = Cost)) +
geom_line(show.legend=FALSE, linewidth = 1) + facet_wrap(Cost ~ ., scales = 'free', ncol = 2) + 
theme_bw() +
theme(plot.title = element_text(hjust = 0.5, size = 10, face = 'bold'), 
panel.grid.major = element_blank(), 
panel.grid.minor = element_blank(),
strip.text.x = element_text(size = 8),
strip.background =element_rect(fill=NA,linewidth=NA),
axis.text = element_text(size = 6)) + 
scale_x_continuous(expand = c(0,0)) +
scale_y_continuous(expand = c(0,0)) +
ggtitle("Cost versus Slope using Canonical Parameters For Various Costs") + ylab("Cost")

pdf("D:/lbmech/Figs/CostFuns.pdf", width = 6, height = 4)
plot(gg)
dev.off()

```

## Velocity plot

```{r velocityPlot}
pdf("D:/lbmech/Figs/VelocityPlot.pdf", width = 6, height = 4)
plotVelocity(velocity_gps)
dev.off()
```

Elevation, ocean currents, and locations:

```{r political_map}

# Import shapefiles for Spain at the Municipal level 
# Crop and aggregate to islands
muns <- geodata::gadm("Spain",level=3, path = "GADM")
muns <- project(muns,proj)
muns <- crop(muns,region)
islands <- aggregate(muns)

# Download coarser DEM for plotting purposes; remove data below sea level
dem <- rast(elevatr::get_elev_raster(raster::raster(project(currents,"+proj=longlat")), z = 9))
dem[dem < 0] <- NA
dem <- project(dem,proj)
dem <- crop(dem,plotreg)

# Calculate slope and aspect for hillshade
slope <- terrain(dem,v='slope',unit='radians')
aspect <- terrain(dem,v='aspect',unit='radians')

# Calculate a hillshade raster
hill <- shade(slope,aspect)
names(hill) <- "shades"

# Get a pallete of greys for hillshading
pal_greys <- hcl.colors(1000, "Grays")

# Round the hillshade values to the nearest factor 
hill <- (hill - minmax(hill)['min',]) / (minmax(hill)['max',] - minmax(hill)['min',])
hill <- ceiling(hill * (length(pal_greys)-1)) + 1

# Get colors for the elevation DEM subplot
grad_hypso <- hypso.colors2(10, "dem_poster")
names(dem) <- "Elevation"

# Transform currents into data.table to turn into vector arrows
field <- project(currents, fix_z(proj,res=8000),align=TRUE)
field <- crop(field,plotreg)
field <- as.data.table(rastToTable(field))
names(field)[1:2] <- c("u","v")
field <- na.omit(field)

# ggplot2 call
ggplot() +
geom_spatraster(data= dem) +    # Elevation is the basemap
scale_fill_gradientn(" \nElevació (m)",
colours = hypso.colors(15,'dem_poster'), na.value='#005EB8', # Color palette
limits = c(0,unlist(global(dem,'max',na.rm=TRUE)))) +    
theme_bw() +                            # Thematic factors
theme(legend.position = "top",
legend.text = element_text(angle = 45, size = 15),
legend.title = element_text(size=20),
strip.text = element_blank(),
strip.background =element_rect(fill=NA,linewidth=NA),
panel.grid.major = element_blank(),
panel.grid.minor = element_blank()
) + 
guides(fill = guide_colorbar(barwidth = 65, barheight = 1.5, alpha = 0.7,
label.position = "top",
override.aes = list(alpha = 0.7))) +
geom_spatraster(data = hill, fill = pal_greys[as.matrix(hill)],  # Hillshade layer
alpha=0.4, 
maxcell = Inf) +
geom_spatvector(data = muns, fill=NA,    # Municipal layer
size=1,col = '#EEEEEE') + 
geom_spatvector(data = islands,     # Islands layer
linewidth = 2.5, col = '#454545',fill=NA) + 
geom_quiver(field, mapping = aes(x=x,y=y,u=u,v=v),center=TRUE, linewidth = 0.75,
col = 'lightblue', inherit.aes=FALSE) +   # Ocean currents layer
geom_spatvector(data = pobles, color = 'red', size = 5) +  # Location layer
ggrepel::geom_label_repel(data = pobles, mapping = aes(x = geom(pobles)[,'x'],
y = geom(pobles)[,'y'],
label = pobles$Location),
size = 7) +
scale_x_continuous(position = 'top', expand=c(0,0)) + 
scale_y_continuous(position = 'right', expand=c(0,0)) + 
annotation_scale(location = 'br', height = unit(1,'cm'),text_cex = 3) + 
annotation_north_arrow(location = 'tl',
style = north_arrow_nautical,
height = unit(4.5,'cm'),
width = unit(4.5,'cm'))
```

Cost raster and path figure

```{r, echo = FALSE,message=FALSE,warning=FALSE, fig.height=11, fig.width=8.5, results= 'hide'}
# Smaller region than before
reg <- as.polygons(ext(buffer(pobles, 10000)),crs = crs(proj))

# Uncomment if you've never run this section before before, since the figure
# differs from the example given in the text for clarity

## Get combinations of all paths to and from Palma de Mallorca
# path <- pobles$Location[pobles$Location != 'Palma de Mallorca']
# path <- c(rbind(rep("Palma de Mallorca",length(path)+1),
# path))
# path <- path[1:(length(path)-1)]

## Calculate the paths
# palma_paths <- getPaths(region = reg,
#                   nodes = pobles,
#                   costname = 'energyCosts',
#                   order = path,
#                   id = 'Location',
#                   dir = rd)

# Import the least-cost paths
palma_paths <- vect(paste0(rd,"/world/costrasters/PalmaPaths.shp"))

# Convert currents to a (smaller) data.table
field <- project(currents, fix_z(proj,res=8000),align=TRUE)
field <- crop(field,reg)
field <- as.data.table(rastToTable(field))
names(field)[1:2] <- c("u","v")
field <- na.omit(field)

# Get even coarser DEM for hillshading
dem <- rast(elevatr::get_elev_raster(raster::raster(project(currents,"+proj=longlat")), z = 8))
dem[dem < 0] <- 0
dem <- project(dem,proj)
dem <- crop(dem,reg)

# Calculate aspect, slope
slope <- terrain(dem,v='slope',unit='radians')
aspect <- terrain(dem,v='aspect',unit='radians')

# Calculate a hillshade raster
hill <- shade(slope,aspect)
names(hill) <- "shades"

# Get a pallete of greys for hillshading
pal_greys <- hcl.colors(1000, "Grays")

# Round the hillshade values to the nearest factor 
hill <- (hill - minmax(hill)['min',]) / (minmax(hill)['max',] - minmax(hill)['min',])
hill <- ceiling(hill * (length(pal_greys)-1)) + 1

# Import the cost rsters
costs <- rast(paste0(rd,"/World/CostRasters/Pobles.tif"))

# Keep only the ones involving Palma
palma <- costs[[stringr::str_detect(names(costs),"Palma de Mallorca")]]

# The time ones
palma_dt <- palma[[stringr::str_detect(names(palma),"dt_")]]

# SpatVector Collection with to, from rasters for time
palma_dt_paths <- svc(palma_paths[stringr::str_detect(palma_paths$segment,"Mallorca_to_") &
palma_paths$cost=='dt',],
palma_paths[stringr::str_detect(palma_paths$segment,"_to_Palma") &
palma_paths$cost=='dt',])

# ggplot call for the from raster with time, involving the islands, hillshade
# cost raster, cost paths, and ocean currents
ggtime_from <- ggplot(islands) + 
geom_spatraster(data = palma_dt[[1]]/3600) +
scale_fill_viridis_c(expression(paste(atop("Hores",
italic("Hours")," (hr)"))),
option='magma',direction = -1,
limits = rowMeans(minmax(palma_dt))/3600) +
geom_spatraster(data = hill, fill = pal_greys[as.matrix(hill)],
alpha=0.4, maxcell = Inf) +
geom_quiver(field, mapping = aes(x=x,y=y,u=u,v=v),center=TRUE,
col = 'white', inherit.aes=FALSE) + 
geom_spatvector(
linewidth = 1.25, col = '#000000',fill=NA) + 
geom_spatraster_contour(data = palma_dt[[1]],binwidth = 3600) + 
geom_spatvector(data = palma_dt_paths[[1]],
linewidth = 1.4, col = 'yellow') +
geom_spatvector(data = pobles,
size = 3, color = 'red') + 
theme_bw() + 
theme(legend.position = "right",
legend.text = element_text(angle = 45, size = 10),
legend.title = element_text(size=12, hjust=0),
axis.title.y = element_blank(),
strip.text.x = element_blank(),
strip.background =element_rect(fill=NA,linewidth=NA),
panel.grid.major = element_blank(),
panel.grid.minor = element_blank()) + 
xlab(expression(paste("Temps des de Palma — ",italic("Time from Palma")))) +
scale_x_continuous(position = 'top', expand=c(0,0)) + 
scale_y_continuous(position = 'right', expand=c(0,0)) + 
ylab("")

# Now likewise for time to
ggtime_to <- ggplot(islands) + 
geom_spatraster(data = palma_dt[[2]]/3600) +
scale_fill_viridis_c(expression(paste(atop("Hores",
italic("Hours")," (hr)"))),
option='magma',direction = -1,
limits = rowMeans(minmax(palma_dt))/3600) +
geom_spatraster(data = hill, fill = pal_greys[as.matrix(hill)],
alpha=0.4, maxcell = Inf) +
geom_quiver(field, mapping = aes(x=x,y=y,u=u,v=v),center=TRUE,
col = 'white', inherit.aes=FALSE) + 
geom_spatvector(
linewidth = 1.25, col = '#000000',fill=NA) + 
geom_spatraster_contour(data = palma_dt[[2]],binwidth = 3600) + 
geom_spatvector(data = palma_dt_paths[[2]],
linewidth = 1.4, col = 'yellow') +
geom_spatvector(data = pobles,
size = 3, color = 'red') + 
theme_bw() + 
theme(legend.position = "none",
legend.text = element_text(angle = 45, size = 10),
legend.title = element_text(size=12, hjust=0),
strip.text.x = element_blank(),
strip.background =element_rect(fill=NA,linewidth=NA),
panel.grid.major = element_blank(),
panel.grid.minor = element_blank(),
)  + 
xlab(expression(paste("Temps cap a Palma — ",italic("Time to Palma")))) +
scale_x_continuous(position = 'top', expand=c(0,0)) + 
scale_y_continuous(position = 'right', expand=c(0,0)) + 
ylab("")


# And likewise for work
palma_dW <- palma[[stringr::str_detect(names(palma),"dW_")]]
palma_dW_paths <- svc(palma_paths[stringr::str_detect(palma_paths$segment,"Mallorca_to_") &
palma_paths$cost=='dW_l',],
palma_paths[stringr::str_detect(palma_paths$segment,"_to_Palma") &
palma_paths$cost=='dW_l',])
ggwork_from <- ggplot(islands) + 
geom_spatraster(data = palma_dW[[1]]/1000/4.184) +
scale_fill_viridis_c(expression("(kcal)"),
option='magma',direction = -1,
limits = rowMeans(minmax(palma_dW))/1000/4.184) +
geom_spatraster(data = hill, fill = pal_greys[as.matrix(hill)],
alpha=0.4, maxcell = Inf) +
geom_quiver(field, mapping = aes(x=x,y=y,u=u,v=v),center=TRUE,
col = 'white', inherit.aes=FALSE) + 
geom_spatvector(
linewidth = 1.25, col = '#000000',fill=NA) + 
geom_spatraster_contour(data = palma_dW[[1]],binwidth =500000*4.184) + 
geom_spatvector(data = palma_dW_paths[[1]],
linewidth = 1.4, col = 'yellow') +
geom_spatvector(data = pobles,
size = 3, color = 'red') + 
theme_bw() + 
theme(legend.position = "right",
legend.text = element_text(angle = 45, size = 10),
legend.title = element_text(size=12, hjust=0),
strip.text.x = element_blank(),
axis.title.y = element_blank(),
strip.background =element_rect(fill=NA,linewidth=NA),
panel.grid.major = element_blank(),
panel.grid.minor = element_blank()
) + ylab("") + 
xlab(expression(paste("Treball des de Palma — ",italic("Work from Palma")))) +
scale_x_continuous(position = 'top', expand=c(0,0)) + 
scale_y_continuous(position = 'right', expand=c(0,0))

ggwork_to <- ggplot(islands) + 
geom_spatraster(data = palma_dW[[2]]/1000/4.184) +
scale_fill_viridis_c(expression("(kcal)"),
option='magma',direction = -1,
limits = rowMeans(minmax(palma_dW))/1000/4.184) +
geom_spatraster(data = hill, fill = pal_greys[as.matrix(hill)],
alpha=0.4, maxcell = Inf) +
geom_quiver(field, mapping = aes(x=x,y=y,u=u,v=v),center=TRUE,
col = 'white', inherit.aes=FALSE) + 
geom_spatvector(
linewidth = 1.25, col = '#000000',fill=NA) + 
geom_spatraster_contour(data = palma_dW[[2]],binwidth =500000*4.184) + 
geom_spatvector(data = palma_dW_paths[[2]],
linewidth = 1.4, col = 'yellow') +
geom_spatvector(data = pobles,
size = 3, color = 'red') + 
theme_bw() + 
theme(legend.position = "none",
legend.text = element_text(angle = 45, size = 10),
legend.title = element_text(size=12, hjust=0),
strip.text.x = element_blank(),
axis.title.y = element_blank(),
strip.background =element_rect(fill=NA,linewidth=NA),
panel.grid.major = element_blank(),
panel.grid.minor = element_blank()
) + ylab("") + 
xlab(expression(paste("Treball cap a Palma — ",italic("Work to Palma")))) +
scale_x_continuous(position = 'top', expand=c(0,0)) + 
scale_y_continuous(position = 'right', expand=c(0,0))

# And finally energy
palma_dE <- palma[[stringr::str_detect(names(palma),"dE_")]]
palma_dE_paths <- svc(palma_paths[stringr::str_detect(palma_paths$segment,"Mallorca_to_") &
palma_paths$cost=='dE_l',],
palma_paths[stringr::str_detect(palma_paths$segment,"_to_Palma") &
palma_paths$cost=='dE_l',])
ggenergy_from <- ggplot(islands) + 
geom_spatraster(data = palma_dE[[1]]/1000/4.184) +
scale_fill_viridis_c(expression("(kcal)"),
option='magma',direction = -1,
limits = rowMeans(minmax(palma_dE))/1000/4.184) +
geom_spatraster(data = hill, fill = pal_greys[as.matrix(hill)],
alpha=0.4, maxcell = Inf) +
geom_quiver(field, mapping = aes(x=x,y=y,u=u,v=v),center=TRUE,
col = 'white', inherit.aes=FALSE) + 
geom_spatvector(
linewidth = 1.25, col = '#000000',fill=NA) + 
geom_spatraster_contour(data = palma_dE[[1]],binwidth =1250000*4.184) + 
geom_spatvector(data = palma_dE_paths[[1]],
linewidth = 1.4, col = 'yellow') +
geom_spatvector(data = pobles,
size = 3, color = 'red') + 
theme_bw() + 
theme(legend.position = "right",
legend.text = element_text(angle = 45, size = 10),
legend.title = element_text(size=12, hjust=0),
strip.text.x = element_blank(),
axis.title.y = element_blank(),
strip.background =element_rect(fill=NA,linewidth=NA),
panel.grid.major = element_blank(),
panel.grid.minor = element_blank()
) + ylab("") + 
xlab(expression(paste("Energia des de Palma — ",italic("Energy from Palma")))) +
scale_x_continuous(position = 'top', expand=c(0,0)) + 
scale_y_continuous(position = 'right', expand=c(0,0))

ggenergy_to <- ggplot(islands) + 
geom_spatraster(data = palma_dE[[2]]/1000/4.184) +
scale_fill_viridis_c(expression("(kcal)"),
option='magma',direction = -1,
limits = rowMeans(minmax(palma_dE))/1000/4.184) +
geom_spatraster(data = hill, fill = pal_greys[as.matrix(hill)],
alpha=0.4, maxcell = Inf) +
geom_quiver(field, mapping = aes(x=x,y=y,u=u,v=v),center=TRUE,
col = 'white', inherit.aes=FALSE) + 
geom_spatvector(
linewidth = 1.25, col = '#000000',fill=NA) + 
geom_spatraster_contour(data = palma_dE[[2]],binwidth =1250000*4.184) + 
geom_spatvector(data = palma_dE_paths[[2]],
linewidth = 1.4, col = 'yellow') +
geom_spatvector(data = pobles,
size = 3, color = 'red') + 
theme_bw() + 
theme(legend.position = "none",
legend.text = element_text(angle = 45, size = 10),
legend.title = element_text(size=12, hjust=0),
strip.text.x = element_blank(),
axis.title.y = element_blank(),
strip.background =element_rect(fill=NA,linewidth=NA),
panel.grid.major = element_blank(),
panel.grid.minor = element_blank()
) + ylab("") + 
xlab(expression(paste("Energia cap a Palma — ",italic("Energy to Palma")))) +
scale_x_continuous(position = 'top', expand=c(0,0)) + 
scale_y_continuous(position = 'right', expand=c(0,0))


```


```{r, echo=FALSE,warning=FALSE,message=FALSE, fig.height=9.5, fig.width=8.5, fig.align='center'}
gg_egg <- egg::ggarrange(ggtime_to,ggtime_from,
ggwork_to,ggwork_from,
ggenergy_to,ggenergy_from,
nrow=3)
# Final figure is the above 6 with ggarrange. Can't do it by facets since the scales are all different. 
pdf("D:/lbmech/Figs/Paths.pdf",width=8.5,height=7.75)
print(gg_egg)
dev.off()

```
````
